# Resonance Raman spectra for the in-situ identification of bacteria strains and their inactivation mechanism

**DOI:** 10.1101/2020.10.22.351163

**Authors:** Dinesh Dhankhar, Anushka Nagpal, Runze Li, Jie Chen, Thomas C. Cesario, Peter M. Rentzepis

## Abstract

The resonance Raman spectra of bacterial carotenoids have been employed to identify bacterial strains and their intensity changes as a function of ultraviolet(UV) radiation dose have been used to differentiate between live and dead bacteria. The enhanced resonance Raman spectra of color-pigmented bacteria were recorded after excitation with visible light diode lasers. In addition, the resonance enhanced Raman spectra enabled us to detect bacteria in water at much lower concentrations (~10^8^ cells/mL) than normally detected spectroscopically. A handheld spectrometer capable of recording resonance Raman spectra in-situ was designed, constructed and was used to record the spectra. In addition to bacteria, the method presented in this paper may also be used to identify fungi, viruses and plants, in-situ, and detect infections within a very short period of time.

## 1. Introduction

Several optical spectroscopic techniques have been used for the detection of bacteria and viruses [1–7]; Absorption spectra of bacteria show an intense band near 270 nm, which has been assigned to protein and DNA/RNA, while the bacterial fluorescence spectra show intense bands, of tryptophan and tyrosine amino acids, in the 350 nm region. We also find that, synchronous fluorescence spectra (Δλ = 65 nm and Δλ = 25 nm) which display two distinct bands at 310 and 345 nm, corresponding to tyrosine and tryptophan amino acids, respectively [6, 8–10].

However, fluorescence and absorption spectra provide a rather limited information; and although fluorescence and synchronous fluorescence can detect very low concentrations of bacteria, the information provided, by these spectra, is limited because many other molecules and bacterial components emit in the same spectral region, that makes the identification of individual molecules difficult. In addition, bacterial components such as DNA do not fluoresce with sufficiently high intensity to be clearly identified, and in addition, DNA spectra are masked by the intense fluorescence of Tryptophan, which is emitted at the same spectral region. An estimate of the concentration of bacteria, is obtained by recording their light scattering intensity at 600 nm (OD600). However, these scattering data provide very little information, concerning bacterial identity and in addition do not differentiate between live and dead bacteria and neither provide information that may be used to identify the bacterial strain.

In contrast, Raman spectroscopy provides incisive information, which allows us to assign bands to specific bacterial component bonds [11–16], therefore, protein compounds such as Tryptophan, Tyrosine and metalloporphyrins, in solution have been extensively studied by Raman spectroscopy [17–19].

Raman spectroscopy, however, has the disadvantage of requiring higher concentration of bacteria for accurate assignment of several relevant (protein, pigments, DNA) weak vibrational bands. For example, for *E.coli* bacteria cells dispersed in water, a conventional Raman spectrum using visible excitation requires ~ 10^12^ cells/mL [20, 21], whereas conventional fluorescence spectra exhibit a much stronger signal even at ~ 10^7^ cells/mL concentrations. This disadvantage, of Raman spectroscopy, can be, partly, overcome by sample concentration (e.g. centrifuging to concentrate the bacteria in a small volume) or by suitable geometries, such as using several meter long hollow fibers filled with bacteria solution. These techniques, are normally complex, and difficult, if not impossible, to use for bacteria detection especially in-situ.

To overcome these disadvantages, of normal Raman spectroscopy, and be able to record sufficiently intense Raman spectra, at the lower bacterial concentrations, needed for identification of bacteria strains, especially in situ, resonance Raman spectroscopy is a promising technique. In this method, the Raman excitation wavelength is tuned to a wavelength near an electronic absorption of the molecule(s) studied. These electronic absorption bands in biomolecules are usually in the ultraviolet (200 nm – 280 nm) where DNA and protein absorb [20, 22, 23]. However, they are also in the visible regions when the bacteria contain pigments such as carotenes and chlorophyll. Ultraviolet resonance Raman (RR) spectra may be rather difficult to record, first, due to the lack of cw lasers emitting at the required UV wavelength of 200nm - 250 nm, in order to avoid Raman bands being overlapped by fluorescence. In addition, for in-situ recording of high-resolution resonance Raman (RR) spectra, the high spectral resolution requirements imply that handheld instruments maybe too cumbersome to be used in the field and further, high UV excitation power laser pulses may damage the sample. These requirements increase the difficulty of recording ultraviolet resonance Raman spectra in-situ, in the field, without special sample preparations.

Therefore, we have recorded resonance Raman spectra excited in the visible region, because it is simpler and has the ability to identify a large number of diverse bacterial strains, whose absorption is in the visible spectral region, and therefore not affected by the difficulties encountered when recording ultraviolet resonance Raman spectra. Visible resonance Raman spectroscopy may also be applied to most, if not all, bacteria that contain color pigments including *Micrococcus luteus* and *Rhodococcus* where carotenoid pigments are abundant and *Serratia marscescene* that contain prodigiosin[24, 25]. In addition to bacteria, the same advantages of resonance Raman spectroscopy, described above, maybe used to record resonance Raman spectra of green plants, fruits, vegetables and other agriculture related species owing to the fact that they also contain visible color pigments such as chlorophyll and carotenes. Fig. S1 displays the absorption spectrum of red colored *Rhodococcus* bacteria that contain blue-green absorbing pigments. Visible resonance Raman techniques may also be used to also detect viruses because it is known that some viruses can induce pigment formation in the host cell [26].

The function of color pigments, which are an integral part of bacteria, and protection from photo-oxidation, by absorbing ultraviolet and visible radiation is thought to be one of their major functions [27–29]. The resonance Raman spectra of these bacteria may be obtained using high performance, readily available inexpensive and compact visible light emitting semiconductor diode lasers. In addition, these diodes can be operated by a small size battery, which is a requirement for the portable & handheld Raman systems, which we have designed, constructed and describe in a later section of this paper.

## 2. Materials and Methods

### Bacteria preparation and Raman spectra recording

*Micrococcus luteus* bacteria were grown on Tryptone Soya Agar (TSA) plates at room temperature. The freshly grown bacteria were removed from the agar plates and suspended in ~ 12mL saline solution by gently shaking it. The bacteria were subsequently centrifuged for 5 minutes at 3300 rpm, using a Fisher Scientific Model 228 centrifuge.

After centrifugation, the supernatant was discarded and the bacteria were re-suspended in fresh saline solution and the centrifugation process was repeated in order to remove the entire remaining agar. The concentration of the final bacterial suspension was determined by their light scattering optical density at 600 nm. The optical densities of the final bacterial suspensions were kept at ~ 0.5 O.D, at 600nm, where the concentration is ~ 10^8^ cells/mL. The bench-top system used for recording the resonance Raman spectra and UV light irradiation has been described by us previously [21]. Briefly, a Horiba Xplora Raman spectrometer with a 100X (0.90 N.A) microscope objective for excitation and collection of the spectra in an 180°-backscattered geometry was used to record the Raman spectra of the bacteria being studied. The excitation, 532nm, 25mW laser light used for generating the enhanced Raman spectra of the carotenoid pigments of *Micrococcus luteus* bacteria. The resonance Raman spectra of the bacteria were recorded as a function of UV irradiation time, 0, 5, 10 and 20 minutes. To record the resonance Raman spectra, 3 mL bacterial suspension was irradiated, in a 1 cm path-length quartz cell, with 250-350 nm, 6mW/cm^2^ light while stirred continually. Before recording the spectra, the samples were centrifuged in order to increase their concentration (effective concentration ~ 10^12^ cells/mL) in order to achieve higher signal to noise ratio. Subsequently, the RR spectra were recorded using 10uL volume of the concentrated bacteria placed on an aluminum mirror. The recorded RR spectra were baseline corrected and their intensity normalized with respect to the lipid band at 1450 cm^−1^, because UV light has minimal effect on this lipid band intensity. The sample were placed on a rather large aluminum mirror for recording the spectra, which also behaved as a heat sink. The temperature at the laser focus spot was measured by two different thermocouple sensors by which a maximum temperature increase from 23 °C to 35 °C was recorded. Further, after every spectrum acquisition, the sample was visually monitored for any signs of damage and no such damage was observed.

In order to determine the enhancement due to resonance excitation, the RR spectra were also recorded before centrifugation (i.e. at lower concentrations of bacteria in water, ~ 10^8^ to 10^9^ cells/mL) using a 10X (0.25 N.A) microscope objective. The recorded spectra showed a relatively low signal to noise ratio owing to the intense Raman bands of water; however even under these conditions, the major carotenoid bands were well resolved and the effects of UV irradiation on these bands and bacteria were clearly displayed in the recorded Raman spectra. Our new designed and constructed handheld system is described later in this paper.

## 3. Results and Discussion

### 3.1 Effect of UV radiation on *M.luteus* bacteria Raman spectra

Fig. 1 shows the resonance Raman spectrum of *Micrococcus luteus* bacteria excited with 532 nm laser light, after the baseline was removed. The most prominent Raman bands at 1525 cm^−1^ and 1155 cm^−1^ are assigned to the carotenoid pigment C=C and C-C bonds stretching vibrations, respectively[30–32], while the intense band at 1002 cm^−1^ is attributed to the C-CH_3_ bond deformation [30]. In addition to these intense bands, we recorded several relatively less intense vibrational bands at ~ 1580 cm^−1^, 1124 cm^−1^ and 745 cm^−1^, assigned to the carotenoid pigment vibrations (Sarcinaxanthin) present in the *Micrococcus luteus* bacteria. The band at 1650 cm^−1^ is assigned to Amide I polypeptide vibrations and the 1450 cm^−1^ band to lipids (H-C-H deformations) [12, 13, 33]. These spectra assignments make it possible to identify and assign the effect of UV radiation on this bacteria strain, other pathogen species and molecules studied (Table S1).

**Fig. 1.**
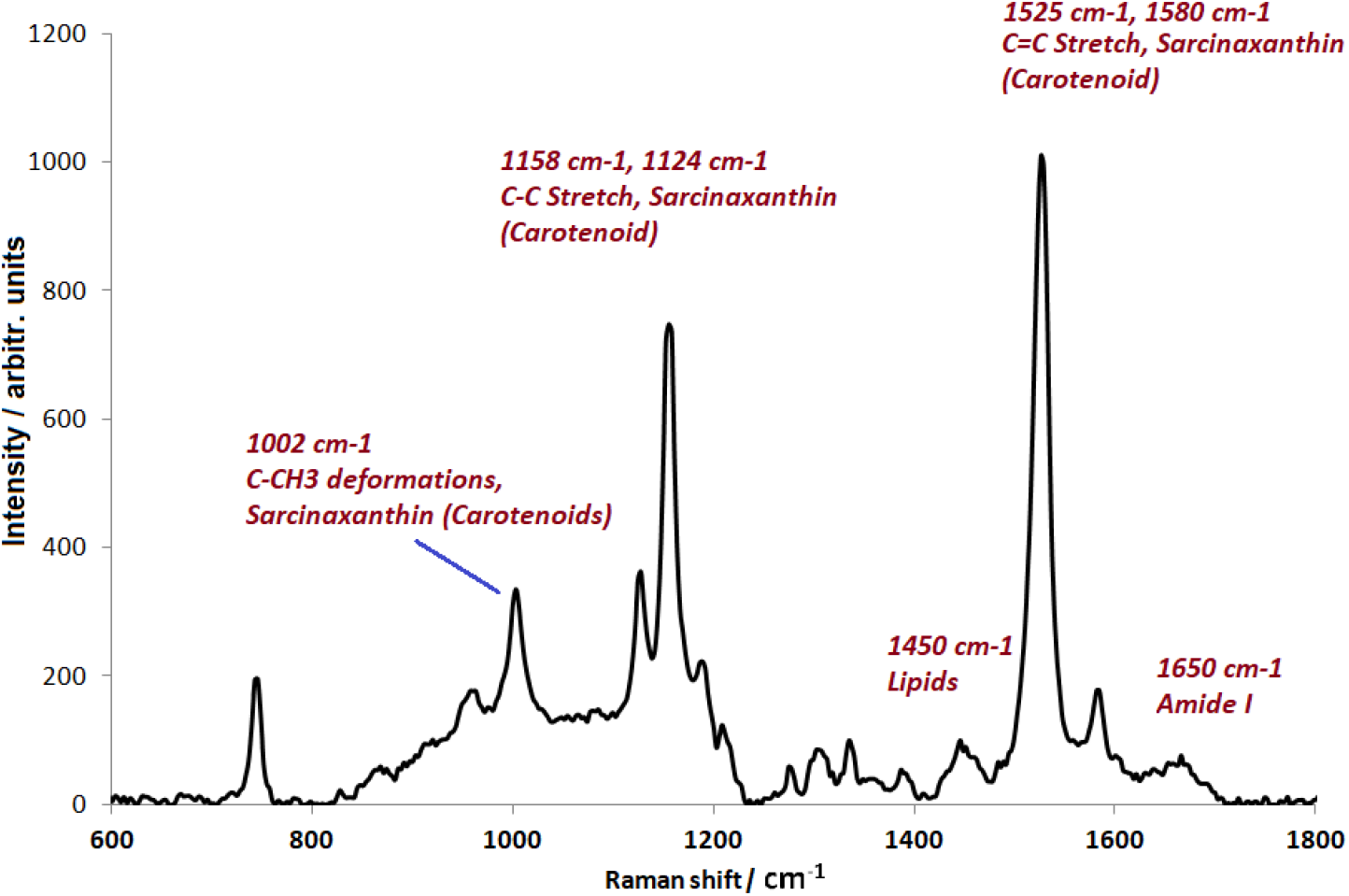
Resonance Raman spectrum of *Micrococcus luteus* bacteria excited with a 532 nm laser. Band assignments are based on [12,14, 31–35].

As expected, owing to the fact that 532 nm laser excitation is very close to the electronic absorption band of the *Micrococcus luteus* bacteria carotenoid pigment (Sarcinaxanthin), a strong enhancement of the Raman bands of the C=C and C-C stretch vibrations is observed.

Fig.2A and 2B show the resonance Raman spectra of *Micrococcus luteus* bacteria as a function of UV light irradiation time, dose. The spectra shown in Fig. 2A are the raw spectra before the removal of baseline. The fitted baseline is also shown. The baseline fit employed was consistent for all the spectra acquired, with 1710 cm^−1^, 1250 cm^−1^, 1065 cm^−1^ and 875 cm^−1^ always on the fitted baseline (See Fig. 2 A). The intensity normalized spectra with respect to 1450 cm^−1^ band after the removal of baseline are presented in Fig 2B. The UV inactivated bacteria are identified by the change in the intensities of their 1525 cm^−1^,1158 cm^−1^, 1127 cm^−1^, 1007 cm^−1^ and 1193 cm^−1^ Raman bands, assigned to carotenoid pigment. It is clearly seen that the ratio between different bands of the bacteria changes as a function of UV irradiation dose. For example, ratio between 1525 cm^−1^ band and 1450 cm^−1^ band, ratio between 1158 cm^−1^ band and 1450 cm^−1^ band or ratio between 1525 cm^−1^ band and 1650 cm^−1^ band etc.

**Fig. 2.**
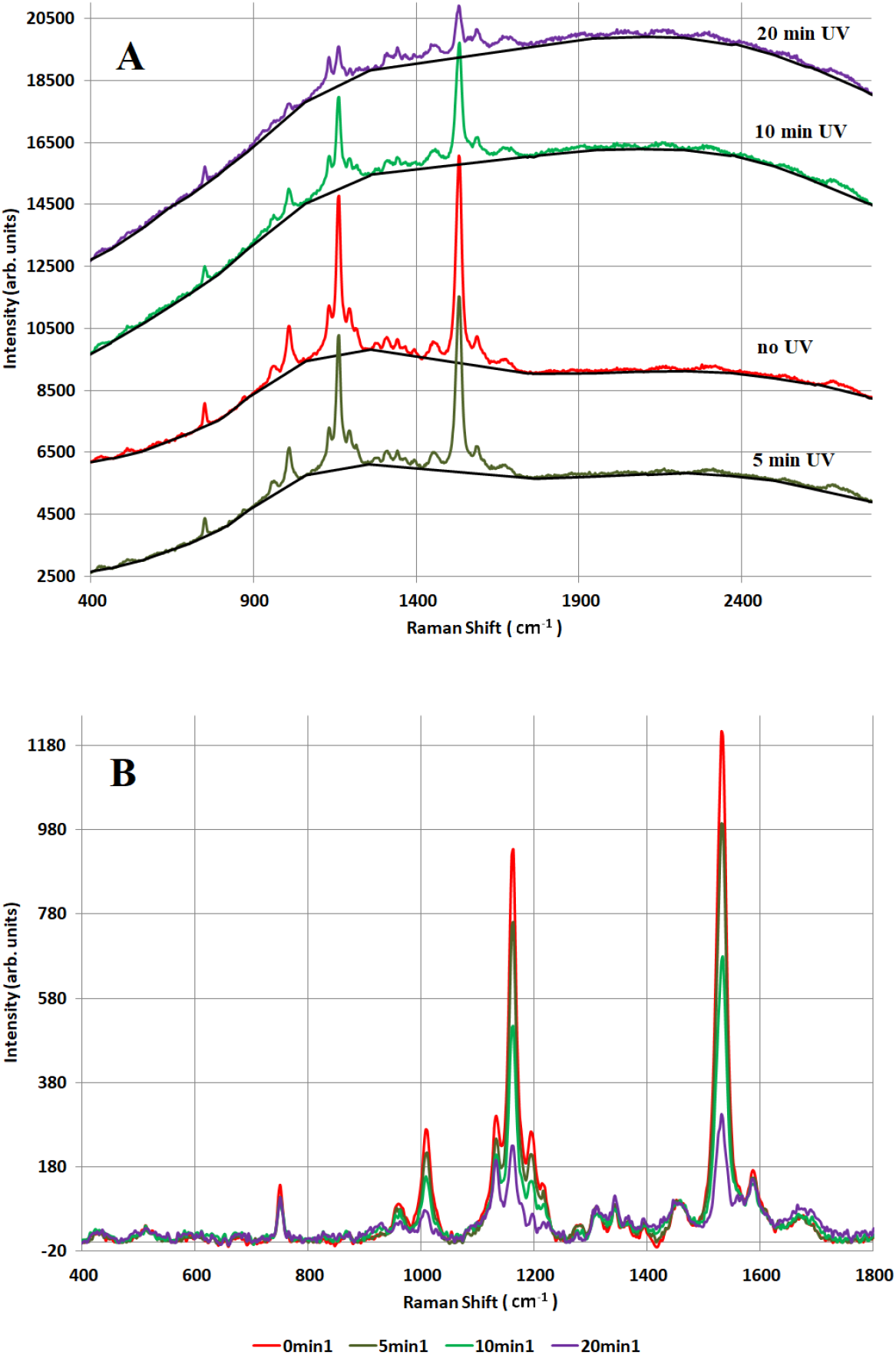
(A) Resonance Raman spectra of *Micrococcus luteus* bacteria changes as a function of UV light irradiation time before baseline correction, the corresponding fitted baselines are shown in black color and (B) Resonance Raman spectra of *Micrococcus luteus* bacteria as a function of UV light irradiation time after baseline correction and normalization with respect to 1450 cm^−1^ lipid band.

We also recorded a rather broad band increase in the intensity of the 1400 cm^−1^ band as a function of UV irradiation time. This increase was seen in several different experimental trials and also with different baseline fits employed. Increase in the same wavenumber region has also been observed by us previously in our studies of *Escherichia coli* bacteria [21]. Therefore this broad increase in the 1400 cm^−1^ region is inherent in the UV irradiated bacteria and is not a part of the baseline. We had earlier assigned this increase to the denaturation of the bacterial proteins due to the formation of UV induced protein photoproducts based on the fact that UV irradiated pure proteins also showed a similar increase in this wavenumber region [21].

Figures 3A and 3B show the ratio of the intensity of the prominent Raman bands, at 1525 cm^−1^ (C=C stretching vibrations of carotenoid polyene) to 1450 cm-1 lipid band and 1158 cm^−1^ (C-C stretching vibrations of carotenoid polyene) intensity to 1450 cm-1 lipid band intensity, respectively, as a function of UV irradiation dose.

**Fig. 3.**
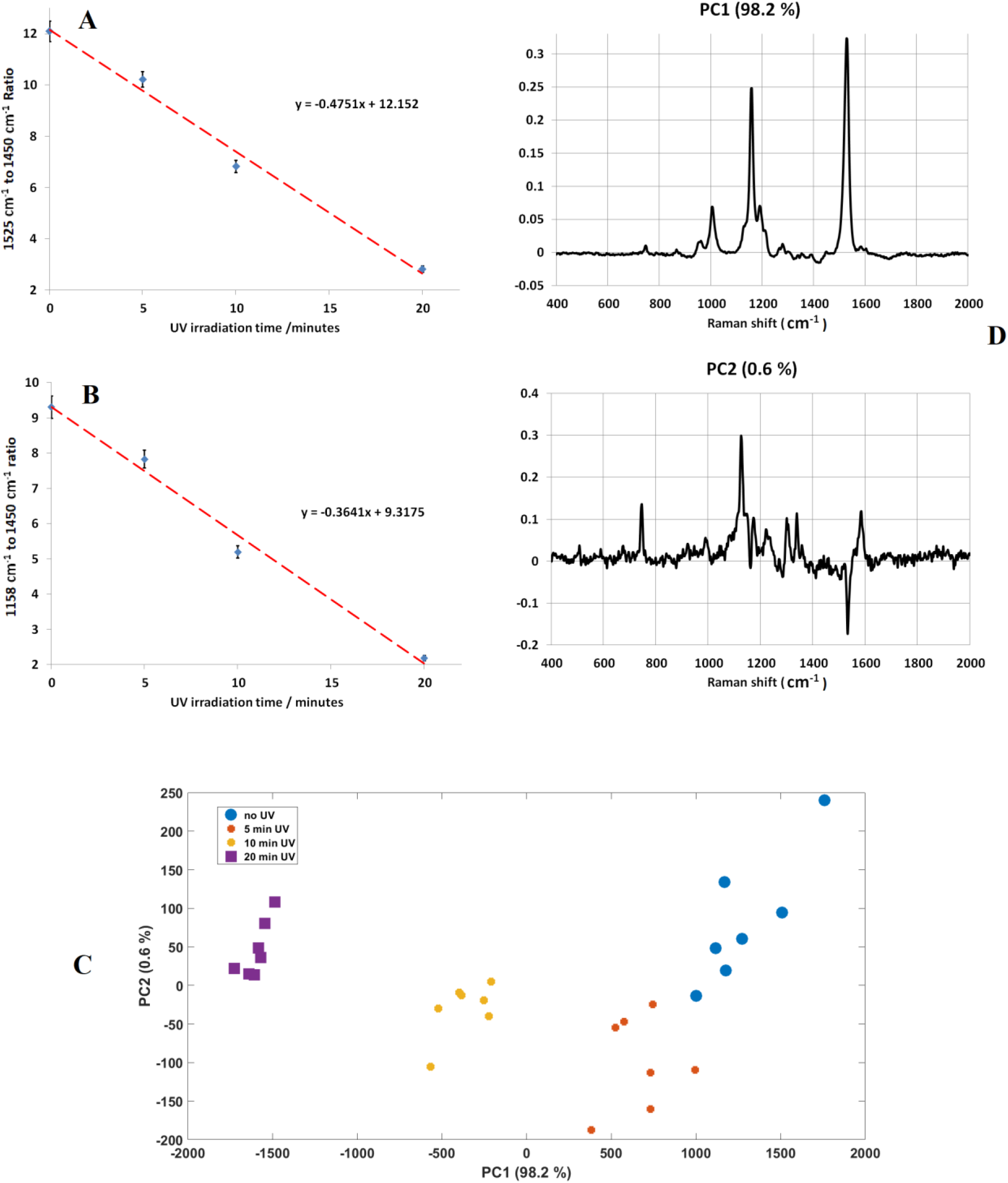
(A) Change in the ratio of 1525 cm^−1^ to 1450 cm^−1^ Raman band intensities of *M. luteus* bacteria as a function of UV dose and (B) Change in the ratio of 1158 cm^−1^ to 1450 cm^−1^ Raman band intensities of *M. luteus* bacteria as a function of UV dose. The error bars represent standard deviation in the values obtained from several different spectra; (C) Principal component analysis plot, from the recorded Raman spectra of *M.luteus* bacteria and (D) plot of Principal component 1 (PC1) obtained after PCA analysis of *M.luteus* bacteria resonance Raman spectra.

We attribute the strong decrease in the intensity of the carotenoid Raman bands with UV irradiation dose to the fact that the carotenoid pigments, in these bacteria have a significant absorption in the ultraviolet region and therefore, the carotenoid pigments bleach during UV irradiation causing the observed decrease in the intensity of their Raman bands. The importance of the assignment of these spectra lies in the fact that they provide the spectral signatures of the UV light inactivated bacteria which we used to quantify the UV irradiation effects on these and other bacteria. Obviously, we do not imply that carotenoids decide the bacteria death, they are simply used as markers. The spectra of the many bacteria components are shown in the figures and their intensity changes are compared to the intensities of the bacterial carotenoids components because the enhanced resonance Raman intensities of the carotenoids allow for a more accurate estimation of the bacteria inactivation as a function of UV radiation and correlated them with the PCA analysis data.

Another significance of result of these experiments is the UV light induced damages to bacteria, which can be correlated with the inactivation of live and formation of dead bacteria. Principal component analysis (PCA) (Fig. 3C) of the recorded Raman spectra depicts a clear separation between the Raman spectra of *M. luteus* bacteria before and after UV inactivation. It is clearly shown in Figure 3C that the UV irradiated spectra move in the general direction of decreasing weight of Principle component 1 (PC1) which accounts for 98.2 % of the variance. Analysis of PC1 (Fig. 5B) shows a pronounced decrease in the Raman bands of carotenoids and a small increase in the 1400 cm^−1^ band, which we assign to protein dissociation products[21]. Therefore, in order to differentiate between live and dead bacteria, the ratio of the prominent carotenoid bands at 1525 cm^−1^ and 1410 cm^−1^ can also be calculated along with the earlier explained ratios of 1525 cm^−1^ to 1450 cm^−1^ band intensity ratios. Plating and subsequent counting of Colony Forming Units (CFU), suggests that, the live bacteria population decreases roughly by 4 to 5 orders of magnitude within 5 minutes of UV irradiation (with UV intensity ~ 5mW/cm^2^). During this time, the ratio of 1525 cm^−1^ to 1450 cm^−1^ ratio, on average, decrease from 12.1 to 10.2, whereas the 1525 cm^−1^ to 1400 cm^−1^ bands ratio, on average, decrease much more sharply due to the fact that these two bands are moving in opposite directions. This sharper change in ratio of 1525cm^−1^ to 1400 cm^−1^ may be able to provide more sensitive measurements for bacterial inactivation for smaller irradiation durations.

It has been shown previously that the carotenoid pigments in bacteria may undergo bleaching under intense laser irradiation [30, 36]. To that effect, in our experiments, we have ascertained that adequate to induce enhanced Raman signal but much less that the required to induce damage to the bacteria.

Fig. 4 depicts the change in the Raman spectra at lower concentrations of *Micrococcus luteus* bacteria after UV irradiation. Under these conditions, the dominant bands located at 3450 cm^−1^ and 1650 cm^−1^are assigned to water molecules. However, even though the water bands are very intense and dominate the Raman spectra of the bacteria recorded, yet, the decrease in the intensity of the prominent carotenoid bands at 1525 cm^−1^, 1158 cm^−1^ are clearly observed. The spectra shown in Fig. 6 were normalized to the 2950 cm^−1^ band after removal of the background baseline.

**Fig. 4.**
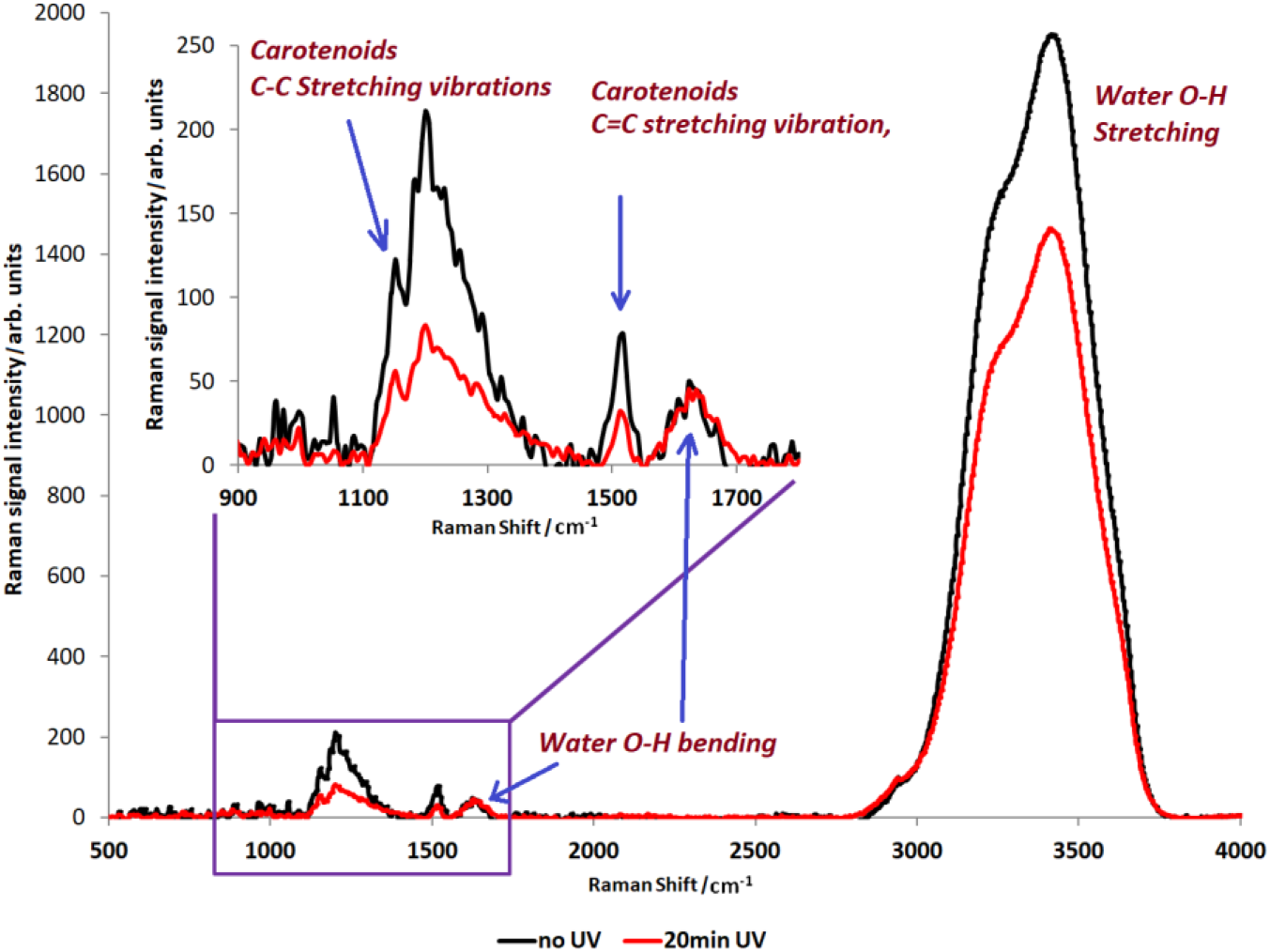
Changes in the Raman spectra of Micrococcus luteus bacteria as a function of UV irradiation. The spectra were recorded at relatively low concentration (10^8^ – 10^9^ cells/mL).

### 3.2 Identification of bacteria strains by means of resonance Raman spectroscopy and Principal Component Analysis (PCA)

It is also possible, to identify different strains of bacteria using resonance Raman spectroscopy, even though many of these bacteria contain similar colored pigments. This is due to fact that the length of the pigment polyene chain, such as carotenoid, affects the frequency at which major vibrational bands appear in the Raman spectra [32]. In addition, the molecules bonded to the chain-ends of the pigment molecules, such as carotenoid, may display unique bands in the Raman spectra, thus making their identification possible. Figure 5A shows the resonance Raman spectra of *M.luteus*, *S. marcescene* and *E.coli* bacteria where the difference in their spectra characteristics is evident. Five spectra were normalized, with respect to the lipid band at 1450 cm^−1^, and averaged after baseline correction. It can be seen in Fig. 5A that *E.coli* bacteria do not show any of the prominent carotenoid bands in their Raman spectra, however, both *S. marcescene* and *M.luteus* show prominent, resonance enhanced pigment bands with unique spectral features, which we have utilized for the identification of pigmented bacteria strains. Fig. 5B shows the Principal Component Analysis (PCA) data derived from a set of Raman spectra, of three different bacteria strains, where the strain is clearly identified in the PCA plot.

**Fig. 5.**
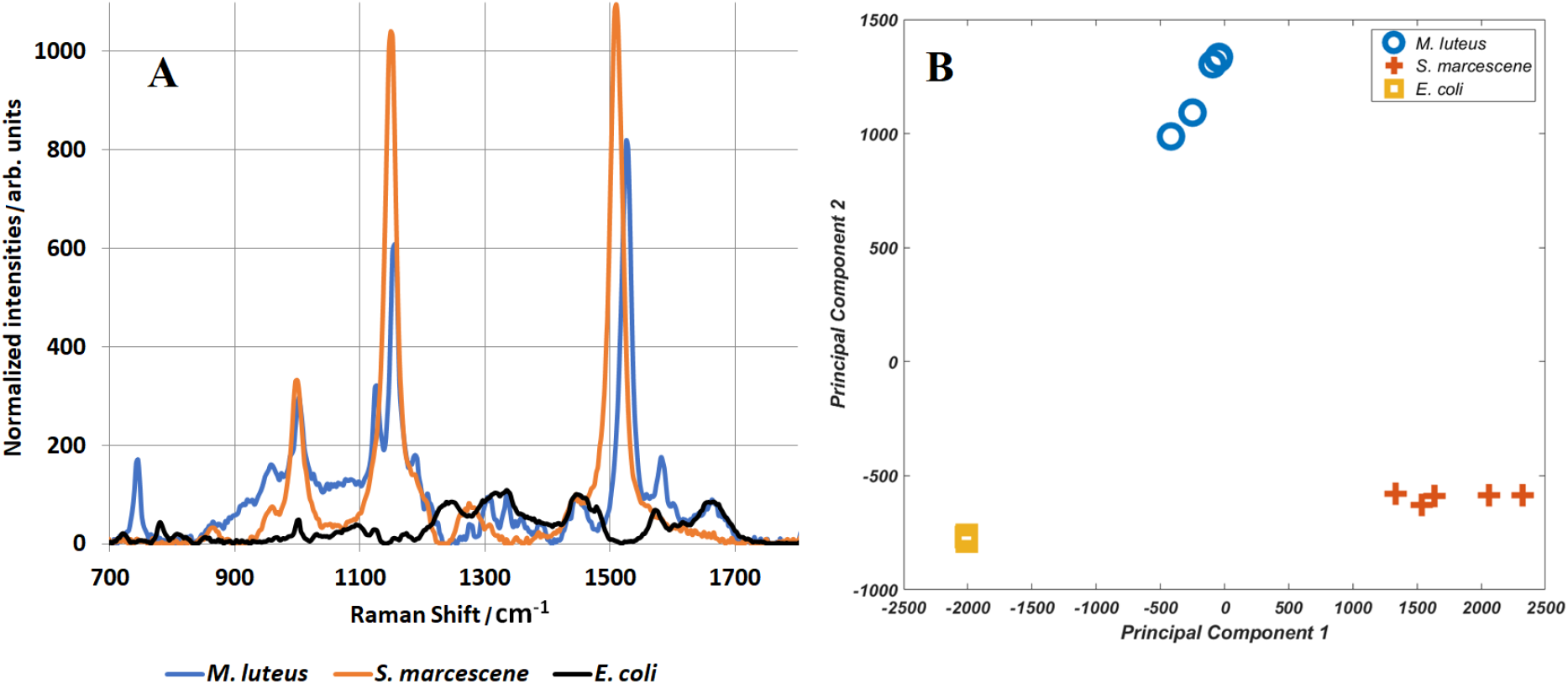
(A) Resonance Raman spectra differences between three bacteria strains.(B) Identification of bacteria strains by applying PCA to the recorded resonance Raman spectra of three different bacteria. The recorded resonance Raman spectra were normalized with respect to the lipid band at 1450 cm^−1^ after baseline correction. The PCA analysis was performed using MATLAB.

### 3.3 Handheld Instruments for in-situ resonance Raman Spectroscopy

The large enhancement of Raman vibration bands by means of visible light resonance excitation induced us to construct a compact handheld Raman spectrometer using easily obtained small size optical components. Our new designed and constructed system is composed of a USB spectrometer with 1800 lines/mm-reflective diffraction grating and a Sony ILX511B linear CCD detector. A diode laser ~50mW, emitting at 532 nm wavelength was used as the excitation source and the spectrometer was calibrated using the Raman spectrum of ethanol as reference. The Raman spectra were acquired using an 180° backscattered geometry (Fig. 6 A), and a 0.25 NA microscope objective which focused the excitation beam on the sample and collected the Raman signal. A dichroic mirror, 532 nm laser line filter, and long-pass filter were used in this handheld RR system were obtained from Thorlabs. This compact (6 inch X 6 inch X 6 inch) system records Raman spectra similar to those recorded by the Horiba bench-top Raman instrument. Fig. 6 B shows the resonance Raman spectrum of *S. marcescene* bacteria recorded by this handheld spectrometer after baseline removal. The spectral resolution of the recorded Raman spectra by the handheld resonance Raman spectrometer is ~ 35 cm^−1^ that is sufficient for bacteria identification.

**Fig 6.**
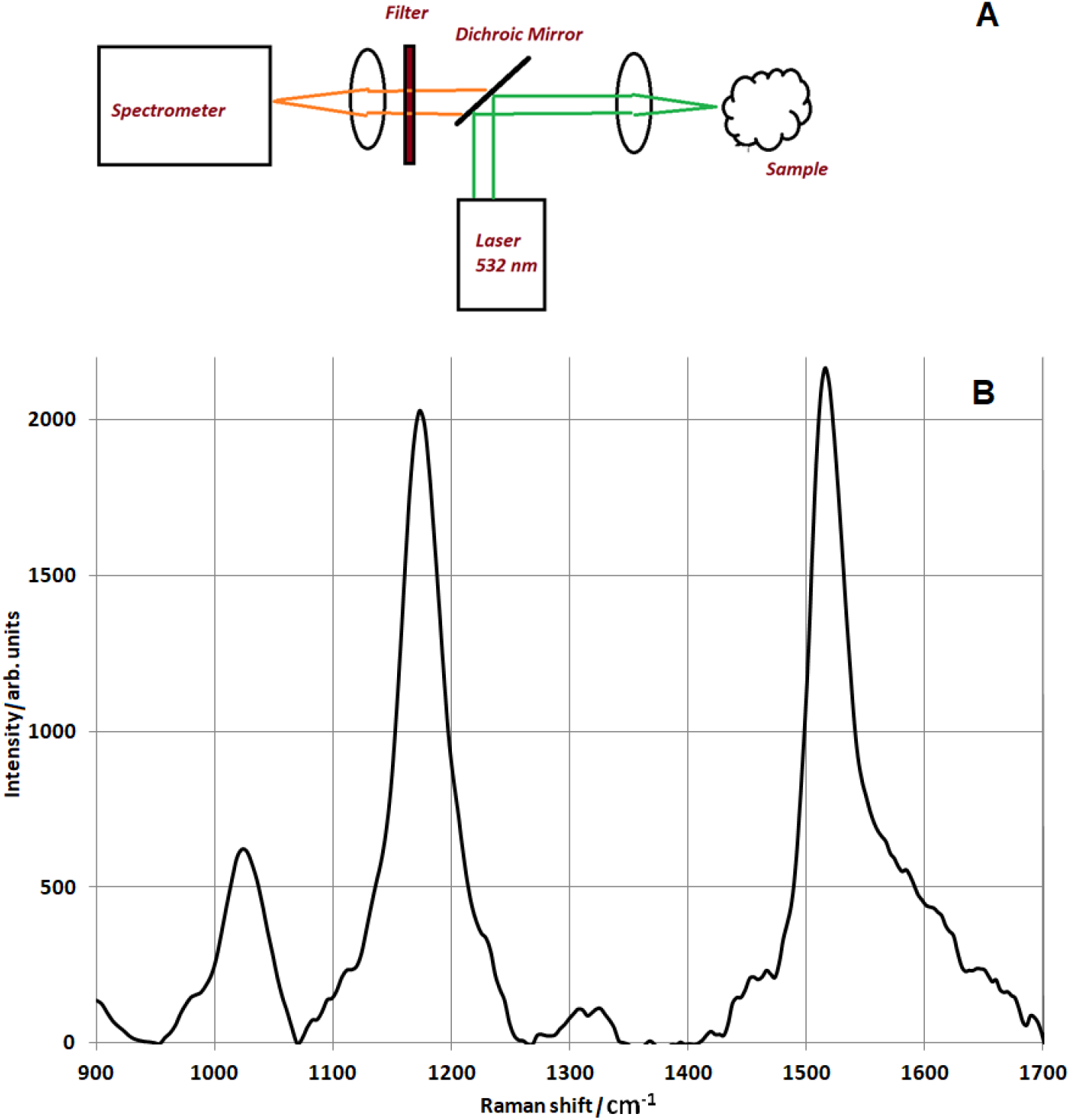
Schematic diagram of the handheld Raman spectrometer system (A). Recorded resonance Raman spectrum of *S. marcescene* bacteria after baseline correction (B).

## 4. Summary and Conclusion

The resonance Raman spectra of color pigmented bacteria, were recorded before and after irradiation with 250nm to 380 nm ultraviolet light, which induces their inactivation. The recorded resonance Raman spectra of live and inactivated (dead) bacteria as a function of UV irradiation dose show prominent changes in the Raman spectra intensities of the carotenoid component bands of the bacteria and the formation of a broad new band at ~ 1400 cm^−1^ assigned to protein photoproducts. These changes were attributed to the dissociation of the colored bacteria carotenoid pigments, the inactivation of live bacteria and the formation of UV induced protein photoproducts. The recorded resonance Raman spectra provide, also, a means for determining the ratio of dead vs live bacteria in-situ as a function of UV irradiation dose. Resonance enhancement of the Raman spectra also enabled us to record the Raman spectra of bacteria in water at low concentrations. A small handheld resonance Raman spectrometer was designed, constructed and used to differentiate between live and UV inactivated (dead) bacteria in-situ within minutes. We show that the recorded resonance Raman bacterial spectra of the hand-held and the benchtop spectrometer are practically identical. The handheld spectrometer may also be utilized to record in-situ, Raman spectra of other biological species such as virus and plants.

The need for the detection of viral pathogens in situ, in addition to bacteria is very important not only as a means for their identification and possible prevention of wide spread diseases but also for the early detection of viral and bacterial infection. The handheld resonance Raman spectrometer described in this paper may also provide a useful means for the detection of bacteria and viruses in human cells in-situ, owing to the fact that some viruses lead to the formation of visible pigments in cells [26].

## Acknowledgments

We are grateful for the partial support of this research by the Welch Foundation Grant 1501928, the Air Force Office of Scientific Research Grant FA9550-20-1-0139, and the Texas A&M Engineering Experiment Station (TEES). The authors would like to thank Prof. Maria King, for providing bacteria samples and valuable discussions and Arjun Krishnamoorthi for technical assistance.

## Supplementary Table and Figures

**Table S I.** Assignment of major Resonance Raman bands in *Micrococcus luteus* bacteria.

| Raman Band (cm<sup>-1</sup>) | Bond Vibrations | Reference |
| --- | --- | --- |
| 1650 cm<sup>-1</sup> | Protein polypeptide backbone<br>vibration | [12, 14] |
| 1525 cm<sup>-1</sup> | Carotenoid, C=C | [31, 32, 34, 35] |
| 1450 cm<sup>-1</sup> | Lipids | [12, 14] |
| 1332 cm<sup>-1</sup> | Amide III vibrations, C<sub>α</sub>-H<br>vibrations, Tryptophan | [14] |
| 1158 cm<sup>-1</sup> | Carotenoid C-C | [31, 32] |
| 1000 cm<sup>-1</sup> | Phenylalanine, Carotenoids, C-C | [12, 31] |
| 745 cm<sup>-1</sup> | DNA, Carotenoids C-C | [12] |

**Fig.S 1.**
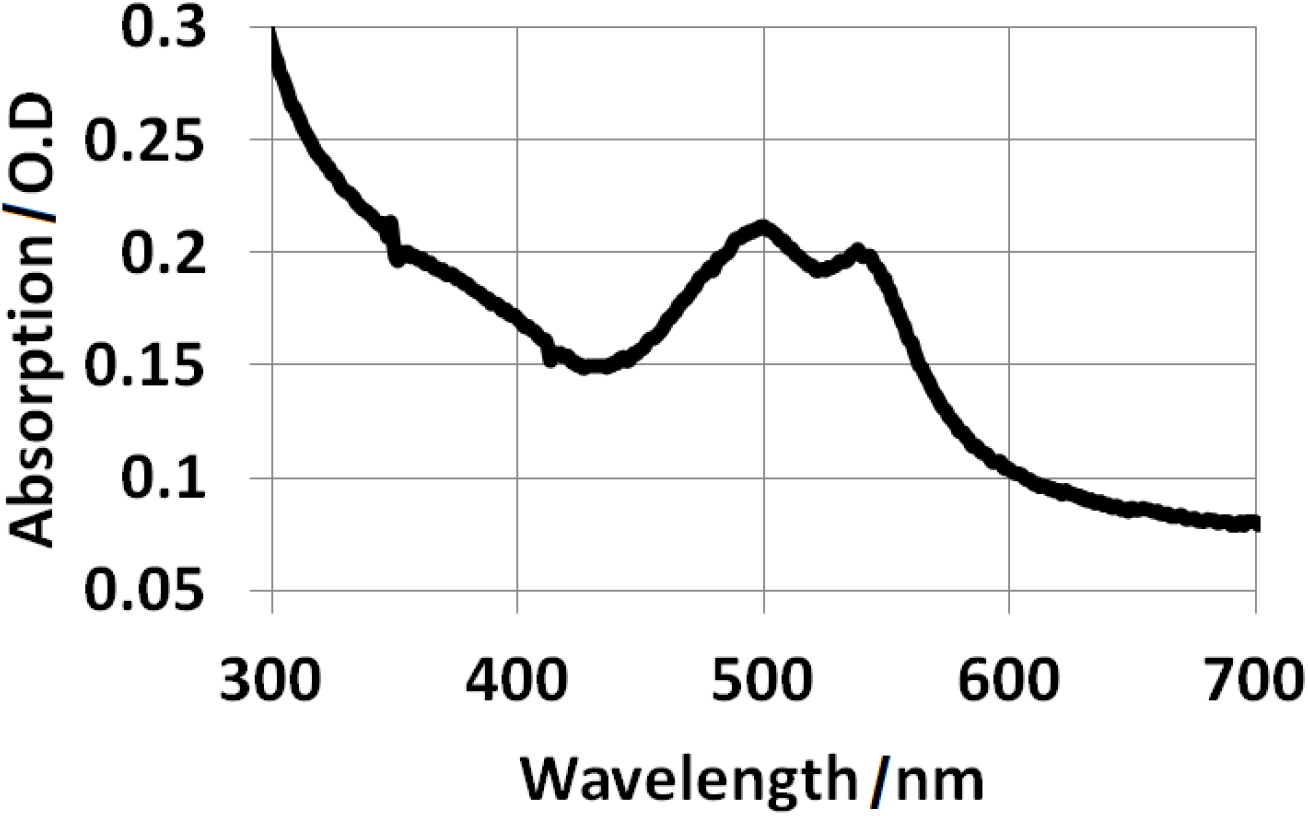
Absorption spectrum of red colored Rhodococcus bacteria; the absorption bands of blue-green absorbing (450 nm – 550 nm) carotenoid pigments are clearly visible.

